# Inhibition of Na/H exchanger-1 in the right ventricle and lung dysfunction induced by experimental pulmonary arterial hypertension in rats

**DOI:** 10.1101/2024.05.27.595780

**Authors:** Giuseppina Milano, Melanie Reinero, Julien Puyal, Piergiorgio Tozzi, Michele Samaja, Florence Porte-Thomé, Maurice Beghetti

## Abstract

**Aims:** Pulmonary arterial hypertension (PAH) is a life-threatening disease that still lacks a direct therapeutic approach targeted to the molecular defects associated with the disease. In this study, we focused on the control of the sodium/hydrogen exchange, which is at the root of impaired regulation of intracellular acidity, as well as of the sodium and calcium intracellular overload. We tested the hypothesis that inhibiting the sodium/hydrogen exchanger isoform 1 (NHE-1) with rimeporide enables the recovery of the pulmonary and right ventricular dysfunction in the Sugen5416/hypoxia rat preclinical model of PAH.

**Methods and Results:** We studied 44 rats divided into two broad groups, control, and Sugen5416/hypoxia. After verifying the insurgence of PAH in the Sugen5416/hypoxia group by transthoracic echocardiography and pulse-wave Doppler analysis, two subgroups were assigned to treatment with either 100 mg/kg/day rimeporide or placebo in drinking water for three weeks. The functional, morphological (fibrosis and hypertrophy) and biochemical (inflammation, signalling pathways) myocardial and pulmonary dysfunctions caused by PAH can be at least partially reverted by treatment with rimeporide. Interestingly, the most striking effects of rimeporide were observed in the right ventricle. Rimeporide was able to improve the hemodynamic variables in the pulmonary circulation and the right ventricle, to decrease right ventricle hypertrophy, pulmonary vascular remodelling, inflammation, and fibrosis. No effect of rimeporide is detected in control rats. We also showed that the protective effect of rimeporide was accompanied by a decrease of the p-Akt/Akt ratio and a stimulation of the autophagy flux mainly in the right ventricle.

**Conclusion:** By specifically inhibiting NHE-1, rimeporide at the selected dosage revealed remarkable anti-PAH effects by preventing functional, morphological, and biochemical deleterious effects of PAH on right ventricle and lung. Rimeporide has to be considered as a potential treatment for PAH.

**Clinical Perspective:** *What is new?:* Pulmonary arterial hypertension (PAH) is a disease with a poor survival despite the progress in therapies, the cause of death remains progressive right ventricular failure. The current treatment are essentially pulmonary vasodilators. An ideal drug would show efficacy in pulmonary vasodilation and remodelling but would also have a direct effect on right ventricular function. - Rimeporide, a sodium/hydrogen exchanger type 1 (NHE-1), decreases right ventricular hypertrophy, pulmonary vascular remodelling, inflammation, and fibrosis.
- Rimeporide is promising as it shows an effect not only on the pulmonary vascular bed but directly on the right ventricle.

*What are the clinical implications?:* By specifically inhibiting NHE-1, rimeporide at the selected dosage revealed remarkable anti-PAH effects by preventing functional, morphological, and biochemical deleterious effects of PAH on right ventricle and lung. - This offers new possibilities of treatment of pulmonary hypertension.
- A direct effect on right ventricular function and remodelling is extremely attractive for diverse forms of pulmonary hypertension.

## 1 Introduction

Pulmonary hypertension, a life-threatening disease with a prevalence of about 1% of the global population ^1^, encompasses a heterogeneous group of conditions characterized by mean pulmonary arterial pressure (mPAP) ≥20 mmHg at rest ^2, 3^. The increased blood pressure in the lung and the right ventricle (RV) causes extra myocardial effort to pump blood through the small circulation leading to RV overload and failure. Of the five classes of pulmonary hypertension based on aetiology ^4^, here we focus on Group 1, or pulmonary arterial hypertension (PAH) with mPAP ≥20mmHg and pulmonary vascular resistance ≥2 Wood Units. Despite its global relevance and the availability of treatments that improve symptoms and prolong life, the age-adjusted mortality rate for PAH patients remains still very high ^5^.

In this study we examine the Na^+^/H^+^ exchanger type 1 (NHE-1), a major contributor to intracellular pH (pHi) regulation ^6^. By causing intracellular Na^+^ accumulation and inducing intracellular Ca^++^ overload via the Na^+^/Ca^++^ exchange, NHE-1 hyperactivation may compromise cell homeostasis. The inhibition of NHE-1 has been shown to be a valuable target for cardioprotection in animal models of ischemia-reperfusion injury, cardiac hypertrophy and Duchenne muscular dystrophy ^7–9^. The beneficial effects of NHE-1 inhibitors in clinical trials performed more than 20 years ago, however, is controversial. While the NHE-1 inhibitor cariporide was effective in the GUARDIAN trial ^10^, important cerebrovascular events were highlighted in the EXPEDITION trial ^11^. Another NHE-1 inhibitor, eniporide, instead revealed ineffective in the ESCAMI trial ^12^.

Rimeporide (RIME), a benzoyl-guanidine derivative, was recently tested to prevent skeletal and cardiac damage in Golden Retriever muscular dystrophic dogs ^13^. By inhibiting NHE-1, RIME limits abnormalities in pHi, intracellular Na^+^ and Ca^++^ preventing Ca^++^ and Na^+^ overload^14^. Granted the status of orphan drug for Duchenne muscular dystrophy by the European Medicines Agency and the Food and Drug Administration, RIME has been tested in a clinical trial ^15^. However, no studies have yet tested NHE-1 inhibitors for the treatment of PAH, except one study examining cariporide in monocrotaline-treated rats, which abrogated the PAH-induced dysfunction through decreased right ventricular NHE-1 expression but without effects on pulmonary intimal wall thickening ^16^.

In the present preclinical study, we aim to test the hypothesis that NHE-1 inhibition by RIME alleviates pulmonary and myocardial dysfunction in a Sugen5416/hypoxia (SuHx) rat model. Although SuHx rats are claimed to recapitulate satisfactorily the clinical findings in human PAH patients, we wanted to validate the reliability of the SuHx model to improve the translatability of these findings to patients. We show that RIME may have substantial curative effects in SuHx rats. This conclusion is supported by several analyses, including myocardial and pulmonary morphology, physiology, hemodynamic and function, as well as targeted biomarkers of fibrosis and inflammation. The recruitment of biochemical pathways related to cellular survival signalling, autophagy and NHE-1 expression is also investigated.

## 2 Materials and methods

This research was performed according to the Swiss Federal guidelines (Ethical Principles and Guidelines for Experiments on Animals) and has been approved by the local Institutional Animal Committee (Service Vétérinaire Cantonal, Lausanne, Switzerland, authorization n. VD 3467). All animal procedures were performed in compliance with the guidelines from Directive 2010/63/EU of the European Parliament on the protection of animals used for scientific purposes. All animals were housed on a 12h/12h light/dark cycle with ad libitum standard chow and water. All efforts were made to minimize animal suffering during the experiments. Original data are deposited in https://zenodo.org/me/uploads?q=&l=list&p=1&s=10&sort=newest.

### 2.1 PAH induction in the Sugen5416/hypoxia (SuHx) model

Figure 1 illustrates the protocol. Adult Sprague-Dawley rats (n=44, initial weight=250-300 g, Charles River, France) were divided into two groups (n=22/each). Control rats were injected with 300 μl dimethyl sulfoxide (DMSO, Sigma Chemicals) and housed for 5 weeks in normal cages. SuHx rats received a single 300 μl dose of SU5416 (20 mg/kg subcutaneous injection, Tocris, Bristol, UK) in DMSO, were placed for 3 weeks in a 10% O_2_ hypoxic chambers and for 2 weeks in room air. Animals were then randomized to RIME or placebo treatment for 3 weeks.

**Figure 1.**
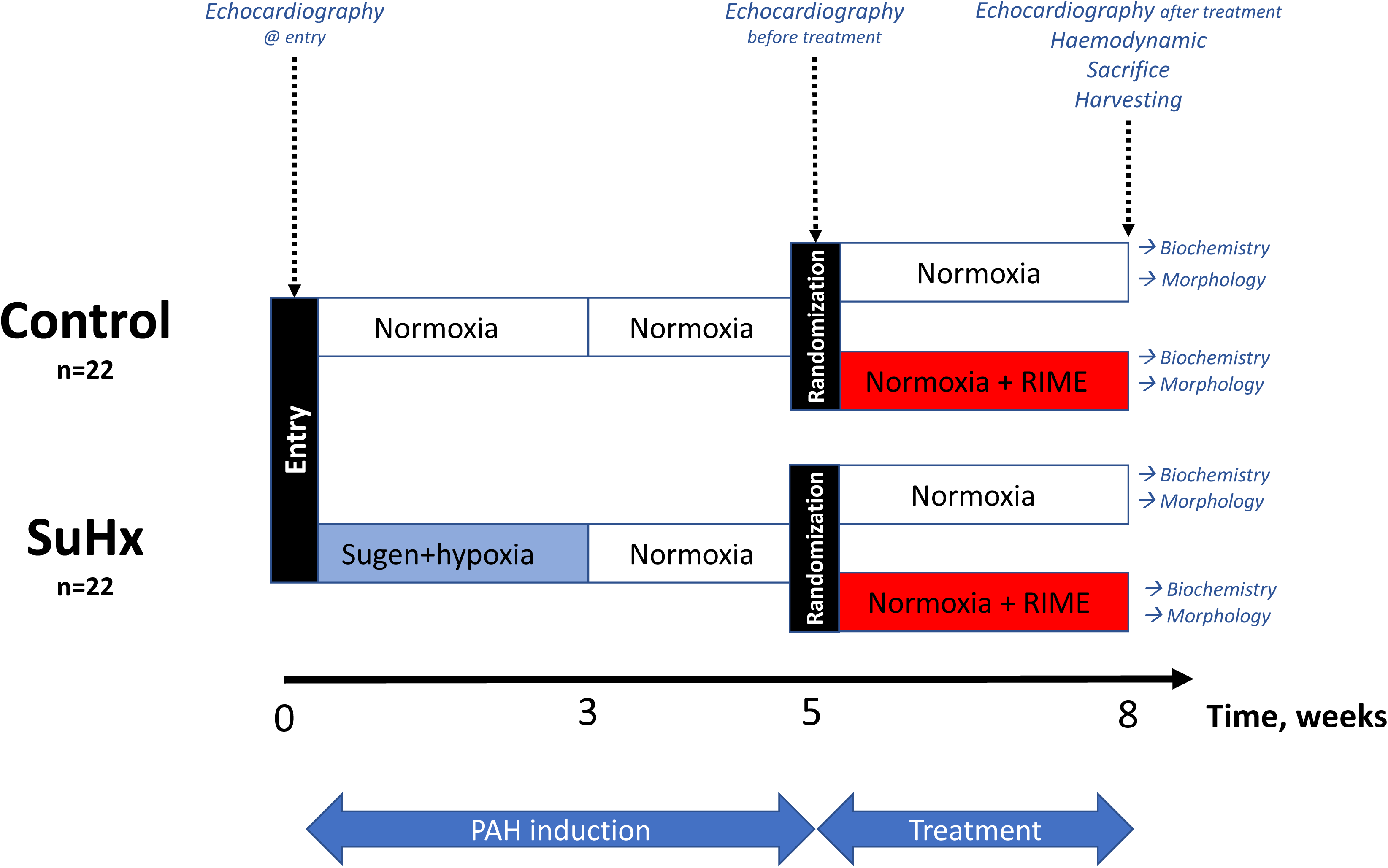
Scheme of the study. In the first part of the study, PAH was induced by injection with Sugen 5416 (20 mg/kg, s.c.) and exposure to hypoxia (10% O_2_) for 3 weeks. In the control group, rats were not injected with Sugen 5416 nor exposed to hypoxia. After randomization, rats were treated either with rimeporide (RIME, 100 mg/kg in drinking water) or with placebo for additional 3 weeks in normoxia. Then rats were assigned to either biochemical or morphological investigations as detailed in the Materials and Methods.

### 2.2 Treatment

RIME powder (EMD-87580) was obtained from EspeRare Foundation (Geneva, CH) as a water-soluble hydrochloride salt. To estimate the quantity dissolved in drinking water, we first determined the average water consumption. Then, solutions containing the equivalent of 100 mg RIME/kg/day in drinking water were prepared daily. At the end of the treatments, rats underwent echocardiography, hemodynamic measurements, blood sampling, and sacrifice.

### 2.3 Echocardiography

All rats were subjected to two-dimension transthoracic echocardiography at study entry, after PAH induction, and before sacrifice using a Sequoia C256 ultrasound machine with a 15 MHz linear array transducer as described elsewhere ^17^. Briefly, anaesthesia was given in an induction chamber with 3% isoflurane. After induction, rats were maintained on isoflurane anaesthesia at 1.0-1.5% supplemented with oxygen delivered at 1.0 L/min and then placed in a supine position on a heated table at 37°C. Right ventricle (RV) and left ventricle (LV) end-diastolic diameter (IDd), end-systolic diameter (IDs) and free wall thickness at diastole (FWTd), were acquired from a parasternal long-axis and analysed offline from M-mode traces at diastole, using Philips Xcelera software. Pulmonary artery (PA) diameter was measured at the level of the pulmonary outflow tract using the superior angulation of the parasternal short-axis view.

Pulsed-wave Doppler was used to measure PA acceleration time (PAT), pulmonary ejection time (ET), and PA velocity time integral (VTI), placing the probe in a parasternal long-axis position and adjusting the angle orientation (in all cases <10°) to optimize visualization of the proximal main PA. mPAP, stroke volume, and cardiac output (CO) were calculated as:

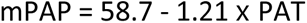

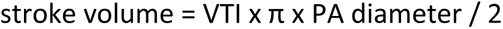

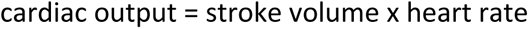

### 2.4 Hemodynamic and arterial blood

At the end of the protocol, after echocardiography, rats anesthetized with 10 mg/kg xylazine, 100 mg/kg ketamine and 1500 IU heparin were placed on a heating pad at 37°C, and ventilated at 50 cycles/min, 2.5 mL tidal volume (Harvard Apparatus model 683, Holliston, MA, USA). The right carotid artery was exposed to withdraw 500 μl arterial blood sample into heparin-coated tubes for blood gas and lactate analysis using i-STAT CG4+Cartridge (Abbott). Then a Millar catheter was introduced into the LV, the 24-gauge needle was connected to MPVS Ultra system (Millar Instruments) to record the LV pressure waves and obtaining end-diastolic pressure (EDP), peak systolic pressure, dP/dt_max_, dP/dt_min,_ and tau (LabChart software). Then the chest cavity was opened, and the Millar catheter was inserted into the RV to measure the RV pressure waves as described ^18^.

### 2.5 Sacrifice

Rats were euthanized with an overdose of 10 mg/kg xylazine and 100 mg/kg ketamine. Lungs and hearts were removed after sternal incision and immediately processed.

### 2.6 RV hypertrophy

RV hypertrophy was assessed through the Fulton’s index ^19^. Briefly, the atria and the major vessels were removed, the RV was dissected away from the LV and the intraventricular septum (LV+S) and both pieces were weighed. RV hypertrophy was assessed by the RV/(LV+S) ratio.

### 2.7 Histological preparation

Lungs and hearts were perfused en-block with PBS via the RV efflux through a small opening in the left atrium and processed for biochemical and histological analyses. For biochemical analyses, lung, RV, and LV tissues were immediately frozen in liquid nitrogen and stored at - 80°C until use. For histological and immuno-fluorescence analyses, the hearts were perfused with 4% paraformaldehyde via the apex while the lungs were inflated with 4% paraformaldehyde at 25 cm water pressure through the trachea for 10 min. Lungs and hearts were excised, post-fixed in 4% paraformaldehyde for 48 h, cryopreserved in OCT medium, and cut into 8-µm sections.

### 2.8 Pulmonary vascular remodelling

The degree of muscularization of pulmonary arterioles was assessed from immunohistochemical staining of the small PA ^18, 20^. Endogenous peroxidase activity was blocked with 3% H_2_O_2_ and 10% methanol in PBS for 10 min. The sections were permeabilized with 0.1% Triton X-100 in PBS for 30 min and blocked with 5% goat serum for 1 h. Sections were incubated with an antibody against smooth muscle α-actin (α-SMA 1:250,

Abcam 5694) overnight at 4 °C, followed by a goat anti-rabbit IgG secondary antibody (1/500, A16096 Invitrogen), developed with 3,3′-diaminobenzidine (DAB, Sigma), and counterstained with haematoxylin and eosin. Transversally cut arterioles were quantitatively analysed at ×40 magnification using the image analysis system Nikon Eclipse 80i camera and NIH image software (Nikon Instruments Inc., Melville, NY, USA). The percent PA thickness was measured by the formula 100 × (perivascular area-luminal area)/luminal area. Ten randomly selected vessels were analysed for each rat in a double-blind fashion.

### 2.9 Fibrosis

RV and LV sections were stained with Picrosirius red for the analysis of fibrosis expressed as percent collagen-positive area in seven randomly selected fields per ventricle. Lung sections were stained for Masson-trichrome (HT15 Trichrome Stain Kit, Sigma) for measurement of collagen-positive area. Analyses were performed using ImageJ software (NIH).

### 2.10 Immunofluorescence

Defrosted cryosection were permeabilized with 0.2% Triton-1x PBS, blocked with 0.1% Triton X-100 and 10% normal goat serum at room temperature for 1 h, and incubated with various primary antibodies. Methodology is outlined in Supplementary data.

### 2.11 Western blot

Protein lysates from lung, RV, and LV were homogenized in RIPA buffer supplemented with protease inhibitors (Roche) as described ^21^ and detailed in Supplementary data.

### 2.12 RNA isolation and quantitative RT-PCR assay

Extraction and purification of RNA from the RV were performed with RNeasy Mini Kit (QIAGEN 74104) and detailed in Supplementary data.

### 2.13 Statistical analysis

Statistical analysis was performed using Prism10 (GraphPad Software, La Jolla, CA). Cardiopulmonary functional data obtained before and after treatment with RIME or placebo, were first tested by full-model two-way ANOVA in the search of interaction between treatment and group. If P<0.05, this test was followed by the Tukey post-test with correction for multiple comparisons aimed at checking significant changes within the same group (i.e., before vs. after treatment) and among the groups. For clarity reasons, the plots do not display the single points but mean±SEM only. Data obtained at the end of the treatments were graphed as violin plots that report every single data point, their distribution, median and quartiles. All the available data points are shown, with no technical replicates. The significance of the differences among the groups was tested by the one-way ANOVA followed, if P<0.05, by the Šídák post-test to adjust the significance level for multiple comparisons.

## 3 Results

### 3.1 PAH induction

Table S2 shows the body and echocardiographic data obtained in rats at study entry and at randomization (t=5 weeks) in controls (non-SuHx) and after PAH induction (SuHx). Marked variations in relevant parameters highlight successful PAH induction. Four rats died before the randomization but none during the treatment with 100 mg RIME/kg/day. Table S3 shows that blood gasses, sO_2,_ BE and lactate were not altered by RIME treatment.

### 3.2 Cardiopulmonary function

**Figure 2A** shows representative transthoracic echocardiography data obtained in either RV or LV at the end of the treatment with placebo or RIME along with averaged data obtained before and after treatment. **Figure 2B** reports averaged data related to the whole heart function. As expected, PAH compromises the myocardial function before RIME treatment, especially in RV. Thus, FWTd, RV EF, stroke volume and cardiac output are less in SuHx than in controls. Lack of difference between Ctrl and Ctrl+RIME shows that the treatment with placebo is ineffective, and that no effect of RIME is evident in non-SuHx rats. In SuHx rats, 3-week treatment with RIME improves stroke volume, FWTd, and cardiac output. In the RV, RIME beneficially affects IDd, and EF with respect to both the value measured before treatment, and the value measured in placebo. Both variables exhibit improvement with respect to the situation before treatment, and to placebo. Some beneficial effects are also observed in the LV, as for improved EF and IDs.

**Figure 2.**
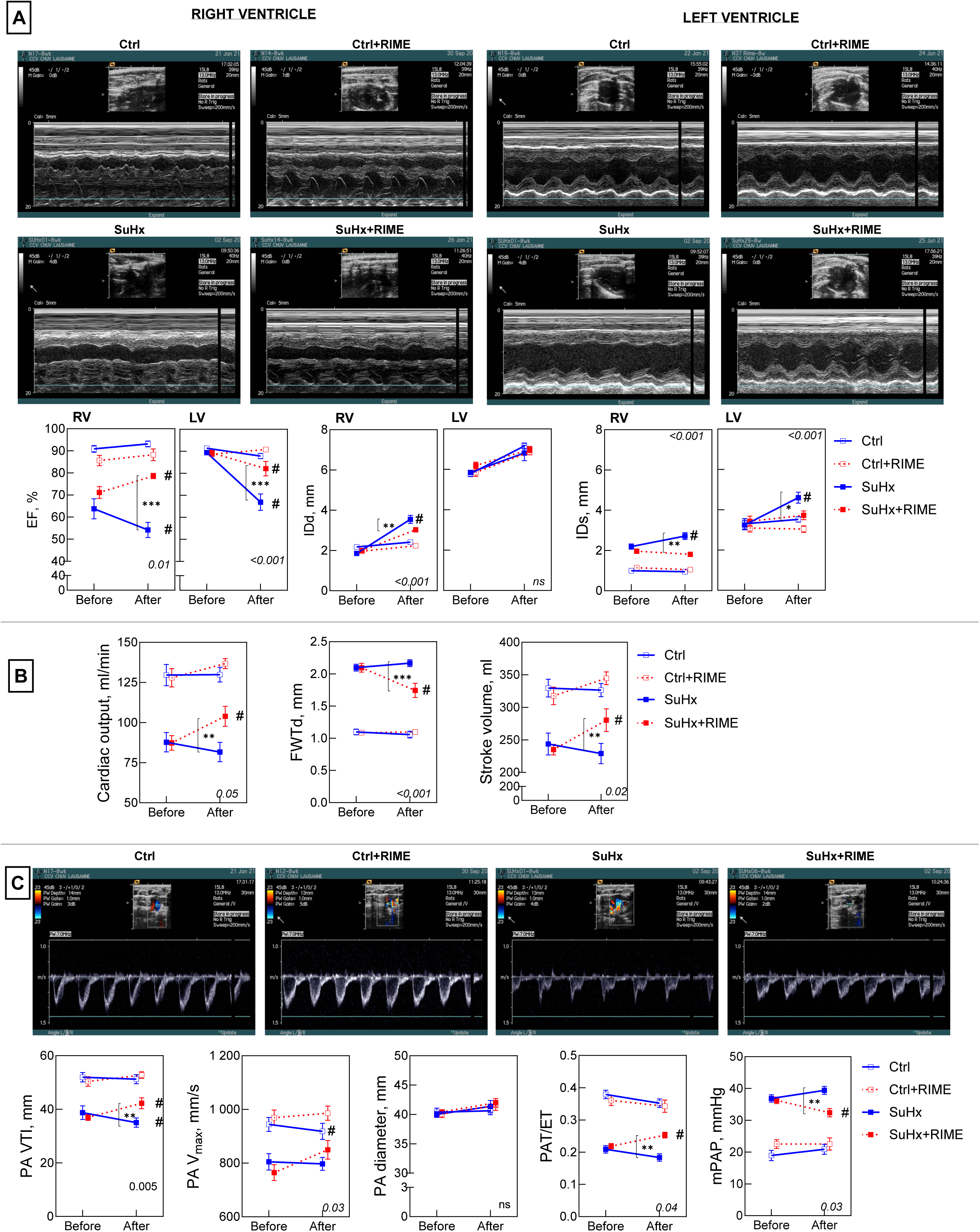
Effect of rimeporide (RIME) on cardiopulmonary function. **Panel A.** Representative M-mode images from RV (left) and LV (right) parasternal long axis at the end of the treatment with RIME. The lower panels display the quantification of the % ejection fraction (EF), internal diameter at diastole (IDd) and systole (IDs). **Panel B.** Cardiac output, free wall thickness at diastole (FWTd), and stroke volume. **Panel C.** Representative transthoracic echocardiography pulsed-wave Doppler flow traces in the pulmonary artery at the end of the treatment with RIME. The lower panels display the quantification of pulmonary artery (PA) velocity time integral (VTI), flow maximal velocity (V_max_), diameter, acceleration time (PAT)/ejection time (ET) ratio, and mean PA pressure (mPAP). Data is obtained before and after RIME in control (empty symbols) and SuHx (filled symbols) rats, treated with placebo (blue continuous lines) or RIME (red dotted lines). Data are expressed as mean±SEM (n=11 animals per group). The two-way ANOVA P values are shown in the inset. If P<0.05, this test was followed by Tukey post-tests to reveal significant changes within the group (before vs. after) or among the groups at the end of the treatment, respectively. # indicates P<0.05 vs. the value measured before the treatment with RIME. ****, P< 0.0001; ***, P<0.001; **, P<0.01; *, P<0.05 when comparing the treatments.

**Figure 2C** shows representative pulsed-wave Doppler data obtained at the end of the treatment with placebo or RIME along with averaged data obtained before and after treatment. As expected, PAH impairs all the markers of the pulmonary function except the PA diameter. Treatment with placebo does not affect any of the markers. No effect of RIME is evident in non-SuHx rats. In the PA of SuHx rats, RIME improves VTI, V_max_, PAT/ET, and mPAP with respect to the values measured before treatment, or the values measured in the placebo group. The improvement associated with VTI and mPAP is evident in both. The PA diameter increases by 2-3% over the value measured before treatment but not significantly. Thus, RIME blocks and sometimes reverts partially the cardiopulmonary dysfunction due to PAH.

### 3.3 Hemodynamic

**Figure 3A** shows representative pressure waves measured in open-chest rats in the RV and LV at the end of the treatment, while **Figure 3B** reports averaged data depicting the hemodynamic. Such data, as well as all those shown in the following figures, was obtained in rats sacrificed at the end of the treatment, thus pre-treatment data is not available. No effect of RIME is evident in non-SuHx control rats. In the LV, PAH does not influence neither the systolic pressure nor dP/dt_max_ but slightly increases dP/dt_min_ and Tau. These pathogenic changes are partially reverted by RIME. By contrast, in the RV, there are marked alterations induced by PAH. RIME exerts protective effects against SP and Tau changes. Thus, RIME partially reverts the RV hemodynamic dysfunction due to PAH.

**Figure 3.**
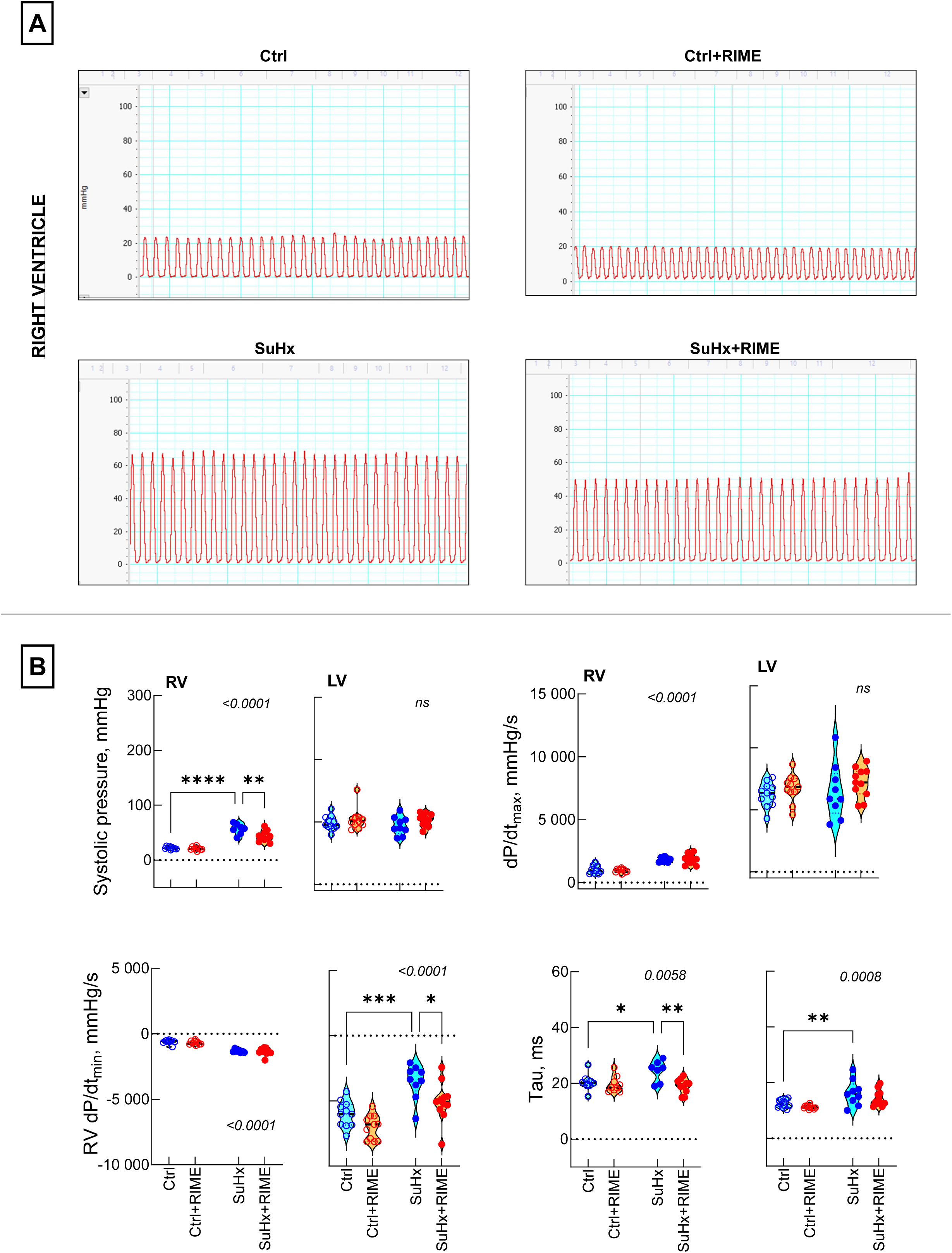
Effects of rimeporide (RIME) on myocardial contractility. **Panel A.** Representative RV pressure traces at the end of the treatment with RIME obtained using a Millar catheter. **Panel B.** Violin plots with distribution, median and quartiles of right (RV) and left (LV) ventricle hemodynamic at the end of the treatments (n=7-10/group). Systolic pressure, maximal rate of pressure development (dP/dt_max_) and relaxation (dP/dt_min_), and tau are shown for each ventricle. The one-way ANOVA P values are shown in the insets. If P<0.05, this test was followed by the Šídák’s multiple comparison test. ****, P< 0.0001; ***, P<0.001; **, P<0.01; *, P<0.05.

### 3.4 Morphology

**Figure 4A**, **4B** and **4C** show representative images obtained in the RV, LV, and lung along with the violin plots obtained from all data. While CSA of cardiomyocytes is an indirect marker of hypertrophy, the pulmonary arterial wall thickness is a marker of lung compliance. All parameters, except the LV cross-sectional area, are impaired in PAH. RIME is effective in reducing such impairment. No effect of RIME is evident in non-SuHx rats. The Fulton’s index is reported in Figure 4A (right bottom panel) as a marker of RV hypertrophy. Thus, RIME partially reverts the morphological dysfunction associated with RV hypertrophy due to PAH.

**Figure 4.**
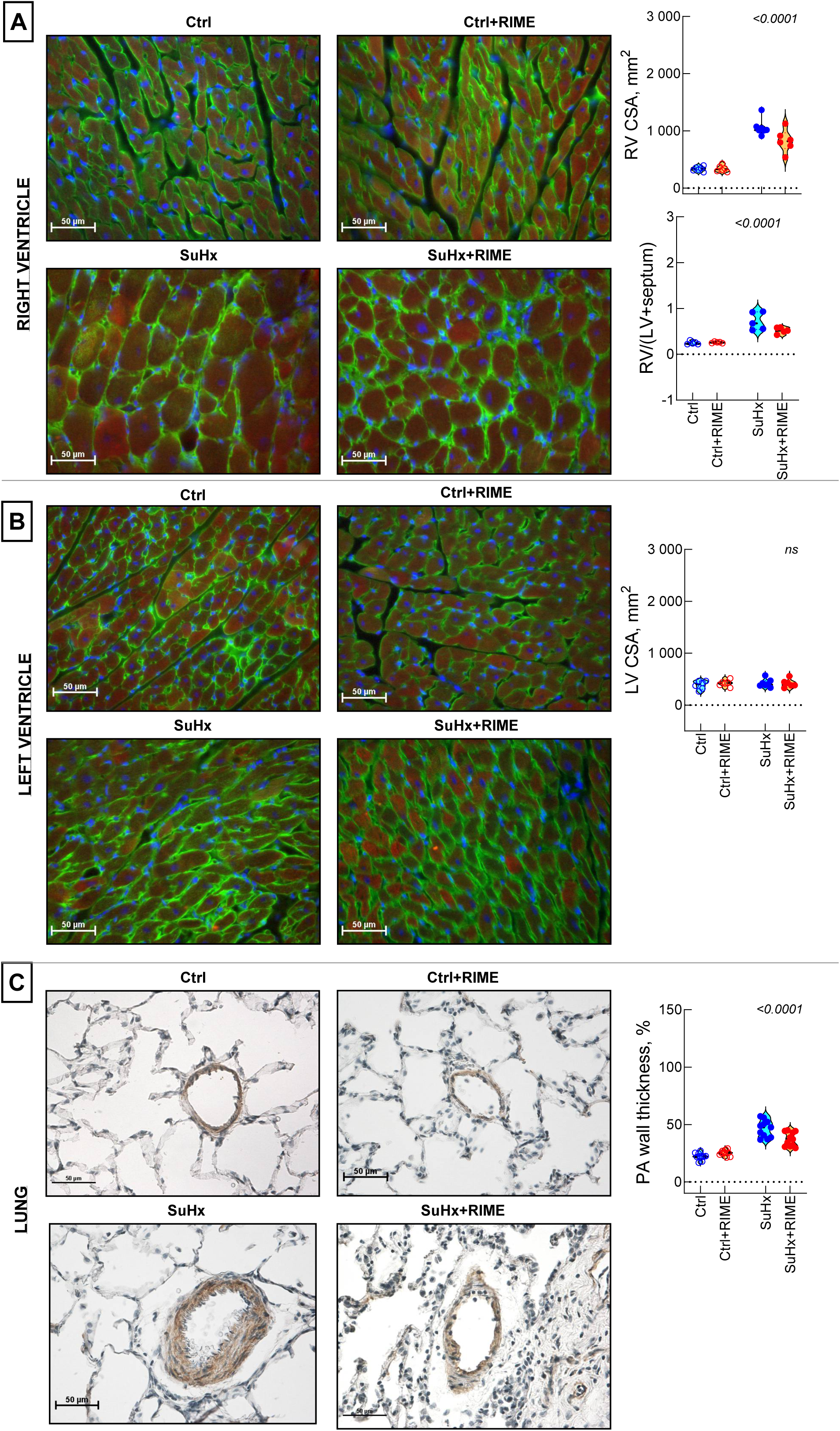
Effects of rimeporide (RIME) on cardiac and pulmonary morphology. **Panel A.** Representative immunofluorescence images of RV obtained at the end of the treatments (n=6-7/group) and stained with antibodies against laminin (green) and α-actinin (red). **Panel B,** same as Panel A but referred to the LV (n=6-7/group). **Panel C.** Representative immunofluorescence images of lung stained with an antibody anti-smooth muscle α-actin at the end of the treatments (n=11/group). The violin plots on the right report distribution, median and quartiles of the cross-sectional area (CSA) (Panels A and B) or pulmonary artery (PA) wall thickness (Panel C) obtained as described in detail in Materials and Methods. Panel A also reports the Fulton index, i.e., the RV/(LV+septum) ratio (n=5/group). The insets show the one-way ANOVA P values. If P<0.05, this test was followed by the Šídák’s multiple comparison test. ****, P< 0.0001; ***, P<0.001; **, P<0.01; *, P<0.05.

### 3.5 Fibrosis

**Figure 5A** and **5B** shows representative images of fibrosis, expressed as collagen-positive areas in cryosections stained with Sirius red in the RV and LV. **Figure 5C** shows representative images of fibrosis in lung as stained with anti-smooth muscle alpha actin (SMA). The right panels of the figures report the respective violin plots. As expected, SuHx groups always exhibit increased collagen-positive areas. In all cases, RIME is effective in reducing fibrosis in SuHx rats. RT-PCR data in **Figure 6D** shows that the RV increase in collagen can be attributed to collagen type I as collagen type III is not affected. Thus, RIME partially reverts the fibrosis associated with PAH, which is almost entirely attributable to increased collagen type I.

**Figure 5.**
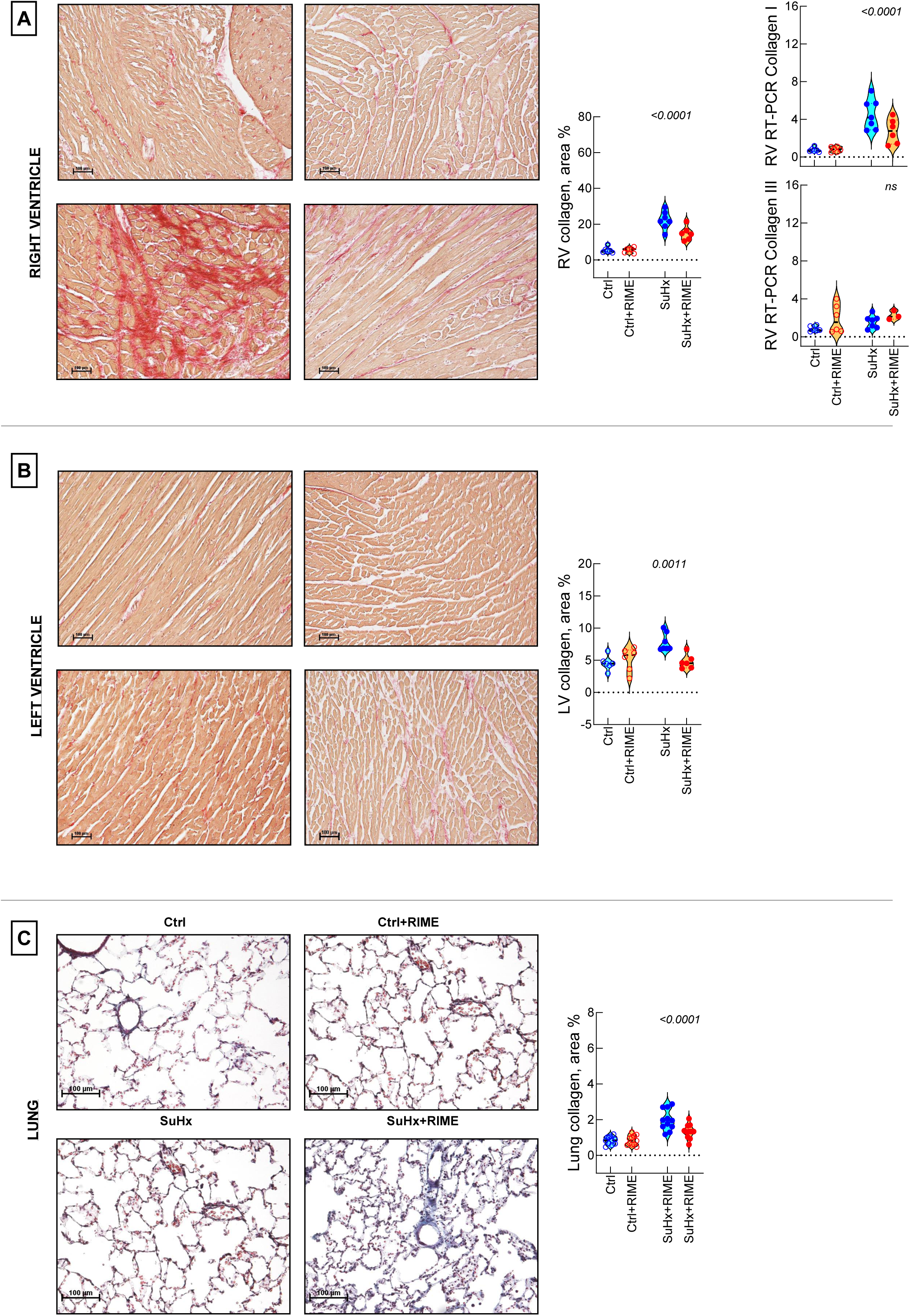
Effect of rimeporide (RIME) on myocardial and pulmonary fibrosis. **Panel A.** Representative immunofluorescence images of RV obtained at the end of the treatments and stained with Picrosirius red, depicting the development of fibrosis (n=6-7/group). **Panel B,** same as Panel A but referred to the LV (n=6-7/group). **Panel C.** Representative immunofluorescence images of lung stained with Masson’s trichome at the end of the treatments (n=11/group). The violin plots on the right report distribution, median and quartiles of collagen-positive area (Panels A and B) or pulmonary collagen (Panel C) obtained as described in detail in Materials and Methods. **Panel D.** Violin plot describing the results of the qRT-PCR analysis of Type I and Type III collagen in the RV (n=6-7/group). The insets show the one-way ANOVA P values. If P<0.05, this test was followed by the Šídák’s multiple comparison test. ****, P< 0.0001; ***, P<0.001; **, P<0.01; *, P<0.05.

**Figure 6.**
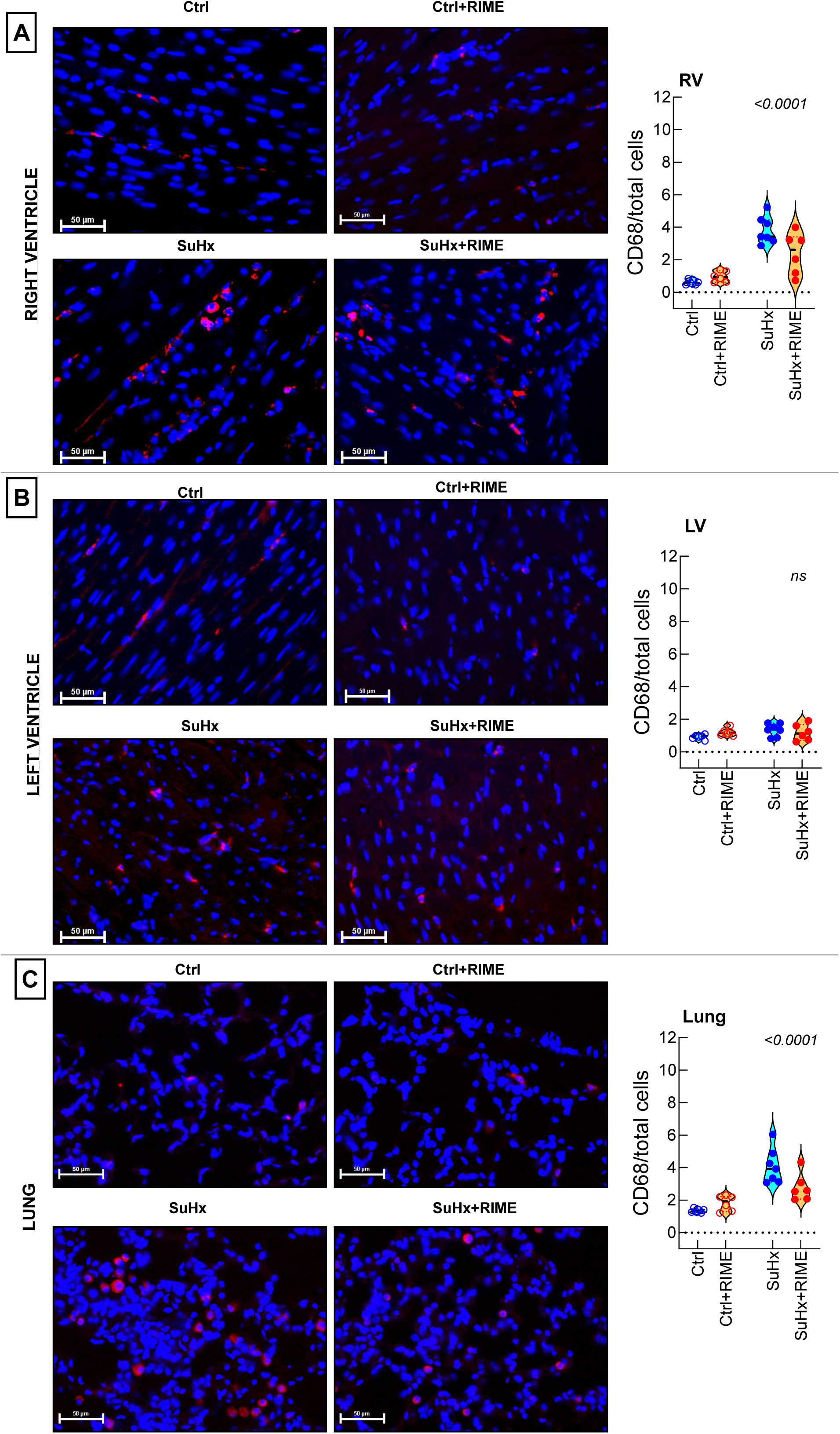
Effect of rimeporide (RIME) on myocardial and pulmonary inflammation. **Panel A.** Representative immunofluorescence images of RV obtained at the end of the treatments and stained with antibodies against CD68 (red) depicting the inflammatory response (n=6-7/group). **Panel B,** same as Panel A but referred to the LV (n=6-7/group). **Panel C,** same as Panel A but referred to the lung (n=6-7/group). The violin plots on the right report distribution, median and quartiles of CD68/total cells obtained as described in detail in Materials and Methods. The insets show the one-way ANOVA P values. If P<0.05, this test was followed by the Šídák’s multiple comparison test. ****, P< 0.0001; ***, P<0.001; **, P<0.01; *, P<0.05.

### 3.6 Inflammation

**Figure 6A**, **6B** and **6C** shows representative images of inflammation, expressed as CD-68 positive cells, in the RV, LV, and lung, respectively. The right panels of the figures report the violin plots derived from all data. PAH induces inflammatory response in the RV and lung, but not in the LV. In both cases, the treatment with RIME significantly reduces CD68, while being ineffective in non-SuHx rats. Thus, RIME partially reverts the inflammation due to PAH.

### 3.7 Signalling pathways

**Figure 7A** reports the expression and the activity of the NHE-1 transporter. PAH increases NHE-1 protein expression in the RV, but not in the lung and LV. In the RV, RIME almost completely blunts the increase observed in placebo. The expression of p-14-3-3 is here assumed to mark NHE-1 activity. As for NHE-1 protein expression, PAH increases p-14-3-3 in the RV, but not in the lung and in LV. Thus, RIME reverts the PAH-induced increase in NHE-1 protein expression and activity.

**Figure 7.**
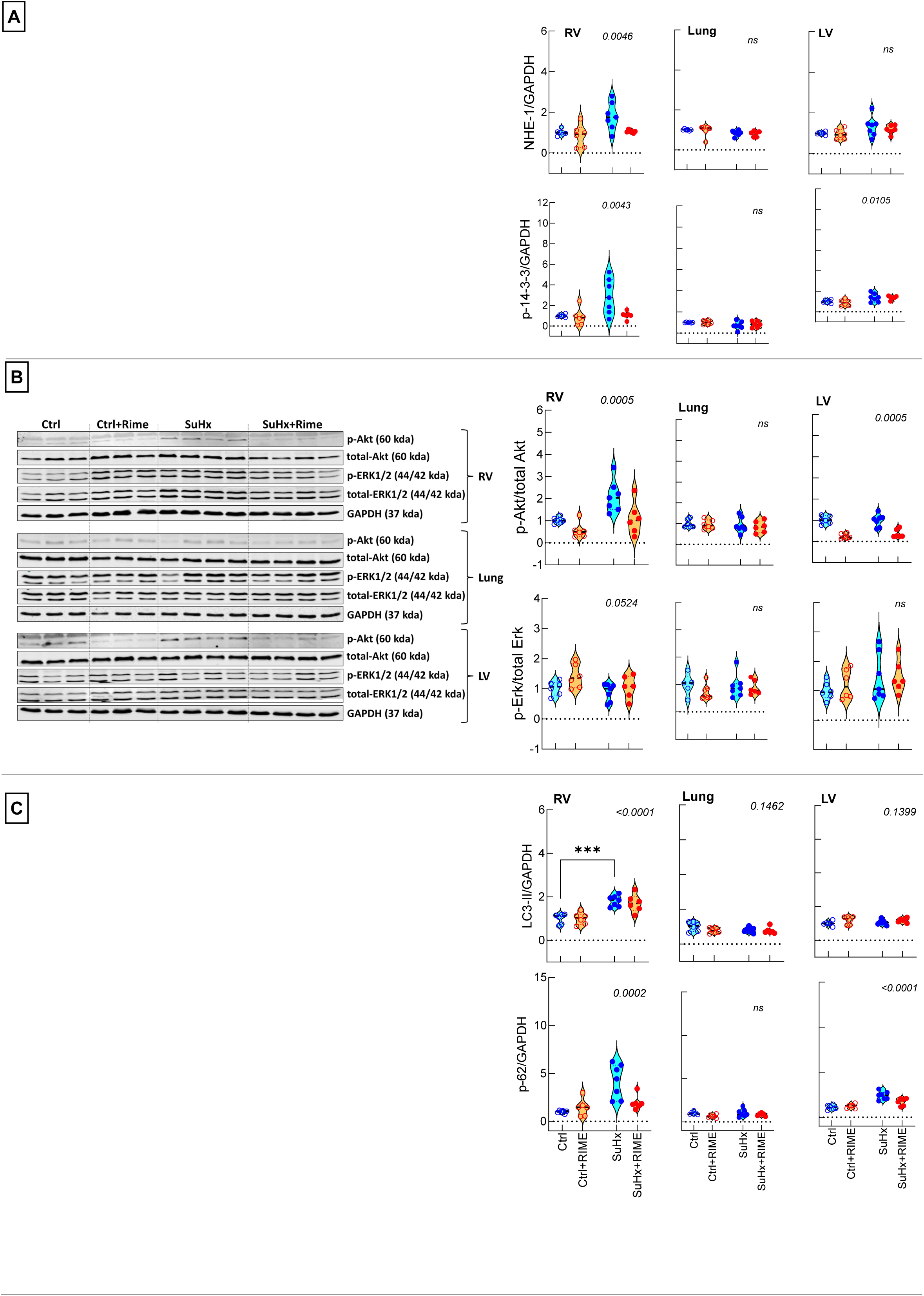
Effect of rimeporide (RIME) on myocardial and pulmonary signalling. **Panel A.** Representative Western blots of RV, lung, and LV of rats at the end of treatments (n=6-7/group). Blots were immunostained against NHE-1 and p-14-3-3 proteins. **Panel B.** Same as Panel A, with blots immunostained against phosphorylated Akt (p-Akt), total Akt, phosphorylated Erk1/2, and total Erk1/2. **Panel C.** Same as Panel A, with blots immunostained against LC3B (upper) and p62/SQSTM1 (lower). Values were normalized relatively to the GAPDH levels. Uncropped unedited blots are reported apart as detailed in the Material and Methods. The violin plots on the right show distribution, median and quartiles of densitometric values obtained as explained in the Materials and Methods section. The one-way ANOVA P value is shown in the insets. If P<0.05, this test was followed by the Šídák’s multiple comparison test. ****, P< 0.0001; ***, P<0.001; **, P<0.01; *, P<0.05.

**Figure 7B** reports the signalling paths associated with Akt and ERK. The most evident changes occur in the RV, where SuHx rats exhibit higher P-Akt/total Akt ratio with respect to control. RIME treatment restores this ratio. In the LV, RIME decreases the P-Akt/total Akt ratio non-significantly in control, but significantly in SuHx rats. By contrast, the ERK pathway seems unaffected by neither PAH nor RIME. Thus, RIME partially reverts the recruitment of the Akt signalling in the RV only, leaving the LV and lung unchanged.

**Figure 7C** reports two main markers of (macro)autophagy. LC3-II and SQSTM1/p62 are markedly upregulated in the RV of SuHx rats. While in the lungs LC3-II does not appear to be changed in the four groups under study, in the LV, a significant increase in SQSTM1/p62 is also detected. In both cases, RIME prevents the SuHx-induced SQSTM1/p62 increases. No significant changes were observed in lung tissue. Thus, RIME partially reverts the upregulation of at least one of the autophagy markers in both the RV and the LV.

## 4 Discussion

In this study, we successfully established a SuHx rat model to mimic PAH disease and showed that PAH can be at least partially reverted by a 3-week treatment with RIME, a potent and selective inhibitor of NHE-1, at 100 mg/kg/day. Remarkably, the effects of RIME are detectable only in SuHx rats. Apparently, RIME targets both the pulmonary vascular remodelling and the RV function, while leaving the LV function and morphology unchanged. The dosage chosen for RIME compares with that tested in the hereditary cardiomyopathic hamster, which prevented the development of necrosis and hypertrophy by downregulating the Na^+^ influx through NHE-1 ^9^.

### 4.1 SuHx model

The Sugen5416/hypoxia model is recognized to closely reproduce the clinical findings in human PAH patients. These include increased precapillary vascular resistance, RV afterload and hypertrophy ^22^, signs of pulmonary vascular lesions similar to the plexiform lesions in human idiopathic PAH ^23^, persistently altered pulmonary vascular remodelling, progressive intima obstruction and high PAP ^24^. Such features lead to decompensated RV failure, elevated RV end-diastolic pressure, RV dilatation and decreased RV EF ^25^, all parameters that were documented to be markedly altered in the SuHx rats used in this study (Table 2). Thus, this model may be considered reliable to test the effects of anti-PAH drugs.

### 4.2 NHE-1 inhibition in PAH

The inhibition of NHE-1 may represent a valuable target to treat PAH as it acts directly on one of its potential causes. Along with the Na^+^/HCO_3_ ^-^ cotransport and the Cl^-^/HCO_3_ ^-^ antiport, NHE-1 is a major regulator of intracellular pH (pHi) ^6^. Human and rat cardiomyocytes preferentially express the NHE-1 isoform ^26^, that controls pHi at the level of the intercalated discs ^27, 28^. Inactive at neutral pH, NHE-1 is activated by acidosis and the myocardial stretch due to hemodynamic overload ^29, 30^. NHE-1 hyperactivation causes intracellular Na^+^ accumulation, which induces Ca^++^ overload via the Na^+^/Ca^++^ exchange, thereby compromising cell homeostasis. NHE-1 inhibition indeed improves cardio-protection in animal models of ischemia-reperfusion and of cardiac hypertrophy ^7–9, 31^.

NHE-1 was upregulated in several models of hypoxia and pulmonary hypertension. In pulmonary arterial smooth muscle cells (PASMC) from chronically hypoxic mice, NHE-1 protein overexpression contributed to PAH development by increasing pHi ^32^. The underlying mechanisms may include the overexpression of hypoxia-inducible factors ^33^ and the acidosis due to enhanced anaerobic glycolysis ^34^. NHE-1 activity was found increased in multiple models of pulmonary hypertension where PASMC proliferation was inhibited by sabiporide ^35^, amiloride ^36^ or selective NHE-1 knockdown ^37^, with the downstream Ca^++^ overload pivotal to increase muscularity and pulmonary vessels contraction ^38^. However, despite solid in vitro and ex-vivo evidence of links between NHE-1 inhibition and PAH, the effects of NHE-1 inhibition on PAH are not yet fully understood. A study highlighted an increase of RV NHE-1 expression in rats where PAH was induced by monocrotaline ^16^. The NHE-1 inhibitor cariporide did not display remarkable effects on pulmonary intimal wall thickening, but nevertheless could attenuate the myocardial dysfunction independently of pulmonary vascular effects ^16^. Thus, the question whether NHE-1 inhibition may target the dysfunction induced by PAH on pulmonary vessels and RV microcirculation remains still needs to be solved.

### 4.3 Rimeporide

The Na^+^/H^+^ exchange is a pleiotropic mechanism that ensures the control of pHi in most mammalian cells ^39^. Of the several diseases potentially triggered by NHE dysfunction, the onset of cardiac fibrosis in Duchenne and Becker muscular dystrophy provided a bench to test RIME as an antagonist of the molecular pathways leading to PAH ^13^. In this model, RIME could limit pHi abnormalities and Na^+^/Ca^++^ imbalance by preventing Ca^++^ and Na^+^ overload ^14^. RIME is presently being tested in a clinical trial that focuses on heart failure prevention in Duchenne muscular dystrophy patients ^15^. The results collected in this study, in facts, converge in highlighting a direct effect of NHE-1 inhibition by RIME in alleviating the outcomes of PAH. Virtually all the parameters affected by PAH tend to return to normal after RIME treatment. Remarkably, RIME affected only the variables related to the pulmonary circulation and the RV leaving those related to the LV nearly unchanged, which highlights a sharply focussed effect targeted at PAH. In this study, we did not assess the activity of NHE-1 but evaluated the expression of RIME protein together with the expression of 14-3-3 binding motif as an indicator of NHE-1 activity. The rationale underlying this choice is based on the finding that the phosphorylation and activation of NHE-1 at Ser703 creates a binding site for 14-3-3 proteins ^40^. This feature was used in an animal model of cardioprotection induced by intermittent hypoxia to monitor the NHE-1 activity though the expression levels of 14-3-3 phosphorylation ^41^. Here we show that PAH increases not only the NHE-1 expression level, but also the expression of 14-3-3 protein. This finding may highlight increased binding of 14-3-3 secondary to increased NHE-1 activity. Both NHE-1 and 14-3-3 protein, however, were downregulated by RIME in the RV, leaving the LV unaffected.

Since the association of autophagy with RV remodelling and failure is poorly studied and because growing evidence suggest that autophagy could play a pivotal role in the development of PAH ^42, 43^, we then decided to evaluate the effect of RIME on autophagy flux in SuHx rats. To monitor the autophagy flux, we investigated the expression of LC3-II (a post-translational lipid-bound modified form of the soluble microtubule-associated protein 1 light chain 3 (LC3-I) which is integrated into the autophagosome membranes) and of p62/SQSTM1 (an autophagy receptor binding to LC3 through its LC3 interaction region ^44^.

Here, we showed that SuHx treatment impairs the autophagy flux in the RV as shown by an accumulation of autophagosomes (increased LC3-II levels) together with a decrease in p62/SQSTM1 degradation. We also observed a slighter effect in the LV (as shown by a slight but significant increase in p62/SQSTM1), suggesting that SuHx is also impairing, but to a lesser extent, autophagy in the LV. We also provide evidence that lung autophagy is not affected. Interestingly, we observed that RIME treatment was able to restore p62/SQSTM1 levels in the heart (both in RV and LV). Since this effect in the RV is associated to increased LC3-II levels (compared to control conditions), RIME treatment appears to boost the autophagy flux in the heart (RV and LV), suggesting that RIME protective effect in cardiac tissue could be associated to autophagy stimulation.

The involvement of autophagy in PAH has been already investigated in the lung but with controversial results. Part of the controversy could be due to the characterization of the autophagy flux in such models/conditions. Because most of the studies have only investigated LC3-II expression which is insufficient to conclude about the flux, some conclusions concerning the involvement of autophagy needs caution.

Lee et al ^45^ demonstrated increased expression of LC3-II in the lung derived both from patients with PAH and mice exposed to chronic hypoxia for 3 weeks. As LC3B(-/-) mice displayed exaggerated PH after chronic hypoxia compared to wild-type mice, LC3B appears to play a highly specific protective role during PH development through downregulation of hypoxic-induced cell proliferation ^45^. Similarly, in the Sugen/hypoxia rat model also the expression of LC3-II in pulmonary endothelial cells increased in parallel with worsening PAH that improved with rapamycin treatment, attenuating RV pressure and pulmonary vascular remodelling ^46^. Other studies performed in monocrotaline rats (another model to induce PAH) have reported contradictory results where the disease progression was associated to increased LC3-II and decreased p62 ^47^. Similar results were observed in human pulmonary artery endothelial cells under chronic hypoxia ^48^. Furthermore, several studies using drugs modulating autophagy (such as rapamycin, chloroquine, and 3-MA) ^46–49^, proposed that autophagy could be a potential novel therapeutical target to treat PAH disease.

Thus, despite the precise mechanisms by which RIME is acting on the autophagy pathway needs more investigation, the present study shows that autophagy is selectively impaired in RV in a preclinical model of PAH and suggests that part of the protective effect of RIME treatment could be associated to stimulation of autophagy.

### 4.4 Clinical implications

Therapeutic strategies available today do not cure PAH, even if they might improve the outcome. The RV function remains the major determinant of poor prognosis and mortality in PAH patients, but so far drug development efforts focused on pulmonary vascular remodelling rather than on heart function in the hope that the RV function improves with improvement in pulmonary hemodynamic. Diuretics, inotropic medications, and, in selected cases, pulmonary vasodilators are indeed empiric medications for managing RV failure by decreasing the filling pressures and afterload rather than drugs that specifically act on the RV function. In facts, none of those therapies significantly improves the long-term outcome in PAH patients, and no targeted RV therapies are yet available, which gives to the results obtained in this study a potential new approach. Trials aimed at testing NHE-1 inhibitors in clinical contexts led to controversial outcomes. While cariporide was successful in patients with unstable angina or non-ST-elevation myocardial infarction undergoing high-risk percutaneous coronary intervention or coronary artery bypass surgery ^10^, important cerebrovascular events were highlighted in bypass surgery patients ^11^, and the NHE-1 inhibitor eniporide was ineffective in patients undergoing thrombolytic therapy or angioplasty for myocardial infarction ^12^. RIME was recently tested to prevent cardiac fibrosis in Duchenne and Becker muscular dystrophy in dogs ^13^. By targeting NHE-1, RIME limits pHi and Na^+^/Ca^++^ imbalance abnormalities and prevents Ca^++^ and Na^+^ overload ^14^. Granted as orphan drug by the European Medicines Agency, RIME is being tested in a clinical trial that focuses on heart failure prevention in Duchenne muscular dystrophy patients ^15^.

### 4.5 Limitations of the study

This is a pilot study that examined one single oral dose of RIME and did not establish if other dosages or routes of administration my provide better delivery of the drug in consideration of its possible use in humans.

Despite we provide evidence that RIME treatment improves the cardiopulmonary function, we did not examine whether these effects of RIME were mediated by the re-establishment of the ox-redox imbalance or by the alleviation of the inflammatory response. As far as the latter is concerned, we believe that the inflammation marker we have used, e.g., the CD68-cells, is a result, not an etiologic agent of PAH.

Finally, we focused only on the NHE-1 isoform without considering, for example, the NHE-8 isoform located in alveolar epithelial cells apical membrane that may greatly influence the Na^+^ transport in alveolar epithelial cells and is under the effect of Ang II in a dose- and time-dependent manner ^50^.

### 4.6 Conclusion

By specifically inhibiting NHE-1 expression and activity, RIME at the selected dosage reveals remarkable anti-PAH effects by reverting partially or totally all the functional, morphological, and biochemical dysfunctions induced by PAH in the SuHx rat model. These effects are mainly targeted to the RV and the pulmonary circulation. These results suggest that RIME could be potentially useful as a drug for the treatment of PAH at the clinical level.

## Acknowledgements

This work was supported by Service agreement between EspeRare Foundation and Centre Hospitalier Universitaire Vaudois (CHUV) and research funding received from Prof MD Maurice Beghetti (Research funds from Cardiology Division University Hospital of Geneva). G.M. and J.P. were supported by a grant from the Faculty of Biology and Medicine of the University of Lausanne. The work on autophagy by J.P. is supported by the Swiss National Foundation (310030L 208141 and 10030_182332).

## Authors’ contribution

Conceptualization: G.M., M.S., F.P.T., M.B.; Methodology: J.P: F.P.T., J-P., G.M., P.G.; Investigation: G.M., M.R., J.P.; Formal Analysis: M.S., G.M.; Resources: F.P.T., M.B.; Writing—original draft: G.M, F.P.T., M.B., J.P.; Writing—review & editing: M.S., G.M., J.P.

## Abbreviations

Akt: protein kinase b
CD68: cluster of differentiation 68
Cq: quantification cycles
CSA: cross-sectional area
DMSO: dimethyl sulfoxide
dp/dt_max_: maximal rate of pressure development
dp/dt_min_: minimal rate of pressure development
EDP: end-diastolic pressure
EDV: end-diastolic volume
EF: ejection fraction
ERK: extracellular-signal regulated kinase
ESV: end-systolic volume
ET: ejection time
ET-1: endothelin-1
FS: fractional shortening
FWTd: free wall thickness at diastole
HR: heart rate
HIF: hypoxia-inducible factors
IDd: internal diameter at diastole
LC 3: microtubule-associated protein 1A/1B-light chain 3
LV: left ventricle
mPAP: mean pulmonary arterial pressure
NHE: sodium/hydrogen exchanger
PAH: pulmonary arterial hypertension
pAkt: phosphorylated Akt
PASMC: pulmonary arterial smooth muscle cells
PAT: pulmonary acceleration time
PBS: phosphate buffered saline
pERK: phosphorylated ERK
pHi: intracellular pH
PSP: peak systolic pressure
RIME: rimeporide
RV: right ventricle
S: septum
SQSTM1/p62: sequestosome 1
SuHx: Sugen5416/hypoxia
Vmax: blood linear velocity
VTI: velocity-integral time.

